# COVIDpro: Database for mining protein dysregulation in patients with COVID-19

**DOI:** 10.1101/2022.09.27.509819

**Authors:** Fangfei Zhang, Augustin Luna, Tingting Tan, Yingdan Chen, Chris Sander, Tiannan Guo

**Affiliations:** Westlake Laboratory of Life Sciences and Biomedicine, Key Laboratory of Structural Biology of Zhejiang Province, School of Life Sciences, Westlake University, Hangzhou, Zhejiang Province, China; Institute of Basic Medical Sciences, Westlake Institute for Advanced Study, Hangzhou, Zhejiang Province, China; Research Center for Industries of the Future, Westlake University, 600 Dunyu Road, Hangzhou, Zhejiang, 310030, China; Center for Infectious Disease Research, Westlake University, 18 Shilongshan Road, Hangzhou 310024, Zhejiang, China; Department of Data Sciences, Dana-Farber Cancer Institute, Boston, MA; Westlake Omics (Hangzhou) Biotechnology Co., Ltd., Hangzhou, Zhejiang Province, China; Systems Biology, Harvard Medical School, Boston, MA

## Abstract

**Background:** The ongoing pandemic of the coronavirus disease 2019 (COVID-19) caused by the severe acute respiratory syndrome coronavirus 2 (SARS-CoV-2) still has limited treatment options partially due to our incomplete understanding of the molecular dysregulations of the COVID-19 patients. We aimed to generate a repository and data analysis tools to examine the modulated proteins underlying COVID-19 patients for the discovery of potential therapeutic targets and diagnostic biomarkers.

**Methods:** We built a web server containing proteomic expression data from COVID-19 patients with a toolset for user-friendly data analysis and visualization. The web resource covers expert-curated proteomic data from COVID-19 patients published before May 2022. The data were collected from ProteomeXchange and from select publications via PubMed searches and aggregated into a comprehensive dataset. Protein expression by disease subgroups across projects was compared by examining differentially expressed proteins. We also visualize differentially expressed pathways and proteins. Moreover, circulating proteins that differentiated severe cases were nominated as predictive biomarkers.

**Findings:** We built and maintain a web server COVIDpro (https://www.guomics.com/covidPro/) containing proteomics data generated by 41 original studies from 32 hospitals worldwide, with data from 3077 patients covering 19 types of clinical specimens, the majority from plasma and sera. 53 protein expression matrices were collected, for a total of 5434 samples and 14,403 unique proteins. Our analyses showed that the lipopolysaccharide-binding protein, as identified in the majority of the studies, was highly expressed in the blood samples of patients with severe disease. A panel of significantly dysregulated proteins was identified to separate patients with severe disease from non-severe disease. Classification of severe disease based on these proteomic signatures on five test sets reached a mean AUC of 0.87 and ACC of 0.80.

**Interpretation:** COVIDpro is an online database with an integrated analysis toolkit. It is a unique and valuable resource for testing hypotheses and identifying proteins or pathways that could be targeted by new treatments of COVID-19 patients.

**Funding:** National Key R&D Program of China: Key PDPM technologies (2021YFA1301602, 2021YFA1301601, 2021YFA1301603), Zhejiang Provincial Natural Science Foundation for Distinguished Young Scholars (LR19C050001), Hangzhou Agriculture and Society Advancement Program (20190101A04), National Natural Science Foundation of China (81972492) and National Science Fund for Young Scholars (21904107), National Resource for Network Biology (NRNB) from the National Institute of General Medical Sciences (NIGMS-P41 GM103504)

**Research in context:** *Evidence before this study:* Although an increasing number of therapies against COVID-19 are being developed, they are still insufficient, especially with the rise of new variants of concern. This is partially due to our incomplete understanding of the disease’s mechanisms. As data have been collected worldwide, several questions are now worth addressing via meta-analyses. Most COVID-19 drugs function by targeting or affecting proteins. Effectiveness and resistance to therapeutics can be effectively assessed via protein measurements. Empowered by mass spectrometry-based proteomics, protein expression has been characterized in a variety of patient specimens, including body fluids (e.g., serum, plasma, urea) and tissue (i.e., formalin-fixed and paraffin-embedded (FFPE)). We expert-curated proteomic expression data from COVID-19 patients published before May 2022, from the largest proteomic data repository ProteomeXhange as well as from literature search engines. Using this resource, a COVID-19 proteome meta-analysis could provide useful insights into the mechanisms of the disease and identify new potential drug targets.

*Added value of this study:* We integrated many published datasets from patients with COVID-19 from 11 nations, with over 3000 patients and more than 5434 proteome measurements. We collected these datasets in an online database, and generated a toolbox to easily explore, analyze, and visualize the data. Next, we used the database and its associated toolbox to identify new proteins of diagnostic and therapeutic value for COVID-19 treatment. In particular, we identified a set of significantly dysregulated proteins for distinguishing severe from non-severe patients using serum samples.

*Implications of all the available evidence:* COVIDpro will support the navigation and analysis of patterns of dysregulated proteins in various COVID-19 clinical specimens for identification and verification of protein biomarkers and potential therapeutic targets.

## Introduction

Since the end of 2019, the world population has been threatened by the severe acute respiratory syndrome coronavirus 2 (SARS2-CoV-2) and the ongoing rise of its constantly evolving variants that have the potential for increased transmissibility, morbidity, and mortality^1^. The spread of coronavirus disease 2019 (COVID-19) shows no signs of being restrained, and drugs with new daily cases worldwide regularly surpassing 1 million^2^. Drugs to treat SARS2-CoV-2 are still insufficiently effective ^3,4^.

Most COVID-19 drugs, if not all, target or act through proteins. Specifically, they mainly target the RNA-dependent RNA polymerase (RdRp) and the main protease of the virus (3CLpro or Mpro), thus inhibiting virus entry and replication^5,6^. However, most of the targeted proteins are not human, partially due to the limited understanding of the molecular dysregulation occurring in patient specimens^7^. Furthermore, proteins are not only relevant as drug targets: they can be robust diagnostic and prognostic biomarkers and the effectiveness of certain drugs can be better assessed via protein measurements.

Using mass spectrometry (MS)-based proteomics, the expression of thousands of proteins can be simultaneously profiled in a variety of patient specimens, including body fluids (e.g., serum, plasma, urine) and tissue (i.e., frozen or formalin-fixed paraffin-embedded (FFPE)). Proteome studies have successfully identified novel biomarkers and drug targets in several clinical studies^8^. Since the first molecular characterization of COVID-19 patient sera^9^, more clinical specimens have been analyzed using mass spectrometry-based proteomics^8,10^. Proteomics data analysis offers unique insights for discovering new potential drug targets. While most published studies analyzed in this research area have focused on the following specimen types: blood samples, including serum^11–21^, plasma ^12,13,22–32^, and peripheral blood mononuclear cells (PBMC)^33,34^, there are also studies analyzing FFPE tissue^35,36^, urine^37–41^, fecal^42^, sputum^23^, extracellular vesicle^43,44^, cerebrospinal fluid^21^, semen^45^, colostrum^46^, colostrum^47^ and nasopharynx swabs samples^48^. All these studies have provided proteomic snapshots of different aspects of tissues from COVID-19 patients. However, few studies have compared the results of multiple studies to fully evaluate this disease due to the lack of proper software tools and databases. While other types of COVID-19 molecular databases exist ^49–52^, none of these is focused on proteomic data from patient samples.

In our meta-analysis, we expert-curated a selection of protein expression datasets published until May 2022, as well as metadata related to the patient and sample information from over 3000 patients. We analyzed the differentially expressed proteins and pathways in various conditions and identifed patterns of recurrently altered protein expression, which can serve as new potential drug targets for treating patients with COVID-19. We also generated a machine learning model for stratifying COVID-19 severity.

## Methods

### Literature search strategy and selection criteria

To produce a comprehensive proteomics data of COVID-19 patients, we used two curation approaches. First, we searched the literature in PubMed using the keywords ‘COVID-19’, ‘patient’, ‘proteomics’, and ‘clinic’. Second, we searched ProteomeXchange, the largest proteomics data repository, using the identifier ‘COVID-19’. Next, we manually went through each study and collected data from their supplementary files. Using these data collation procedures, we thus identified and collected data from 41 studies containing protein expression datasets of COVID-19 patients.

The datasets were organized into tables with patient and sample information, together with the protein expression data. The patient information table includes gender, age, and severity of COVID-19, if available in the original studies. The sample information table describes the types of clinical specimen, the sample preparation, and the methods used for the proteomics data acquisition. For studies using more than two types of clinical specimens, we divided the sample information of each type into separate datasets to facilitate meta-analytic comparisons. The protein expressions were then represented as the measured signals of each protein in each sample. Since protein group quantifications can be ambiguous, we included unique proteins.

## Data analysis

### Patient, sample, and data information

When stratifying patients for analysis, we focused on the information that most studies provide about patients: gender, age, and disease subgroup. The disease subgroups describe each patient's severity level by symptoms. We included the following subgroups: healthy donors, non-COVID-19 controls, COVID-19 (non-severe), COVID-19 (severe), COVID-19 (critical), COVID-19 (non-critical) and COVID-19 (fatal) patients. Disease severity was determined using World Health Organization scores^16^. For several studies, we further classified patients according to their diagnosis of pulmonary fibrosis or their levels of interleukin-6 (IL-6). The datasets were derived from 12 types of clinical specimens: plasma, serum, urine, peripheral blood mononuclear (PBMC), bronchoalveolar lavage fluid, colostrum, extracellular vesicle (EV), feces, nasopharynx swabs, sputa, and FFPE samples derived from heart, kidney, liver, lung, spleen, testis, and thyroid. The sample preparation methods included serum depletion, serum non-depletion, plasma depletion, plasma non-depletion, breast pump, fecal boiling, filter 3kDa, iST kit, methanol precipitation, immune affinity purification, dithiothreitol, ethanol precipitation, acetone precipitation, pressure cycling technology (PCT), RapiGest, red blood cell (RBC) removal, sonication, ultracentrifugation, and others. The proteomics data acquisition methods included data-dependent acquisition (DDA), tandem-mass tags (TMT), enzyme-linked immunosorbent assay (ELISA), multiple reaction monitoring (MRM), data-independent acquisition (DIA), sequential window acquisition of all theoretical fragment ion spectra (SWATH), scanning SWATH (sSWATH), and O-link assays. The proteins included in the database are identified by their UniProt names or HUGO Gene Nomenclature Committee gene names.

### Detection of proteins in different datasets and functional roles

Proteins that were identified in multiple datasets were used for further exploration. We list the fraction of missed detection by mass spectrometry for each protein in each dataset, computed as the percentage of missed detections across all sample files in that dataset. Next, we focused on the 76 proteins that were identified in more than 70% of datasets. The number of unique proteins in the datasets are also listed. The proteins that were consistently identified were analyzed using gene set enrichment analysis using GO, the R databases org.Hs.eg.db^53^, and the package clusterProfiler^54^ for biological process analysis.

### Boxplot analysis of selected proteins

The distributions of the protein abundances were organized by disease subgroups. Specifically, we used grouped boxplots for each dataset and the R package ggboxplot. Unpaired two-sided t-tests were performed with *p*-values calculated by comparing subgroup pairs with the function of t_test in the R package rstatix with a normal distribution assumption. The significance levels of the differential changes were indicated by the corrected *p*-value of 0.5, 0.1, 0.01, and 0.001. For the dataset of Nie *et al.*, the protein expression fold changes were calculated for different tissue types. For Tang *et al.*, Fisher *et al.*, Lam *et al.*, and Zhong *et al.*, which contained information on the disease course, we provided the temporal grouped boxplots with the loess smooth curve fitting regions. For the D'Alessandro *et al.* dataset, the comparative groups were based on the levels of IL-6.

### Pathway analysis of differentially expressed proteins

The molecular pathways containing proteins detected by a specific dataset can be visualized using network graphs and the R package cyjShiny^55^. A pair of disease subgroups can be chosen so that the node size is proportional to the protein expression fold change; the fold change is calculated as the ratio between the mean expression values in each group with an unpaired two-sided t-test. Colors highlight only the nodes with significant changes. The significantly dysregulated proteins were those with a *p*-value < 0.05. KEGG and GO gene set enrichment analyses were performed for such dysregulated proteins using the R package clusterProfiler.

### Co-regulated differentially expressed proteins

Given any two disease subgroups, we identified the proteins that were differentially expressed in the same direction. Fold changes were calculated as the ratios of the mean expression values, the *p-*values were calculated using an unpaired two-sided t-test between the two chosen disease subtypes. The differentially expressed proteins were identified by the user's cutoff fold change and *p*-value. Using a set of sera samples as an example, we identified the proteins that were either up- or down-regulated (adjusted *p*-value < 0.05) in more than five datasets in patients with severe disease vs. patients with non-severe diseases. 51 differentially expressed proteins were used to build a preliminary random forest model to classify COVID-19 severity using the Shen_1 data set as the training set. The resulting top nine proteins were used to build a random forest-based classifier validated in five independent datasets.

### Statistical packages

The statistical analyses of this study used several R packages. Their names and associated version numbers are: org.Hs.eg.db 3.12.0, AnnotationDbi 1.52.0, IRanges 2.24.1, S4Vectors 0.28.1, Biobase 2.50.0, clusterProfiler 3.18.1, cyjShiny 1.0.19, base64enc 0.1-3, graph 1.68.0, BiocGenerics 0.36.1, ggbeeswarm 0.6.0, pheatmap 1.0.12, rstatix 0.7.0, ggpubr 0.4.0, ECharts2Shiny 0.2.13, jsonlite 1.7.2, igraph 1.2.6, htmlwidgets 1.5.3, leaflet 2.0.4.1, shiny 1.6.0, shinydashboard 0.7.1, DT 0.18, plotly 4.9.4.1, ggplot2 3.3.5, shinyWidgets 0.6.0, shinythemes 1.2.0, RColorBrewer 1.1-2, and BiocManager 1.30.16.

## Results

A preliminary set of 41 studies was identified after systematic collection of proteomic data for COVID-19 patients from a set of 316 search results collected from publications as the result of PubMed searches and 178 collected from ProteomeXchange. After manually scrutinizing the full-text and supplementary files of these studies, we selected those containing patient proteomics data for further meta-analyses. The selected studies were further grouped according to their clinical sample type. If a study contained multiple clinical sample types, each sample type was considered a different project with a different dataset. As a result, we collected 53 datasets involving samples from 3,077 patients, 5434 clinical specimens, and 14,403 unique proteins (Figure 1A). For ease of presentation, the projects are represented by author names and a numeric index. We uploaded all 53 datasets to a freely accessible database: COVIDpro (https://www.guomics.com/covidPro/). Using this database, users may select their projects of interest and query for perturbed proteins from specific COVID-19 specimen types. For ease of presentation, the projects are referred to by their first-listed author and PubMed Unique Identifier (PMID). The details of each project are summarized in Appendix 1 and Figure 1B, including the hospital name, city, and nation, as well as the sample type, the MS method, the sample preparation method, and the PMID.

**Figure 1.**
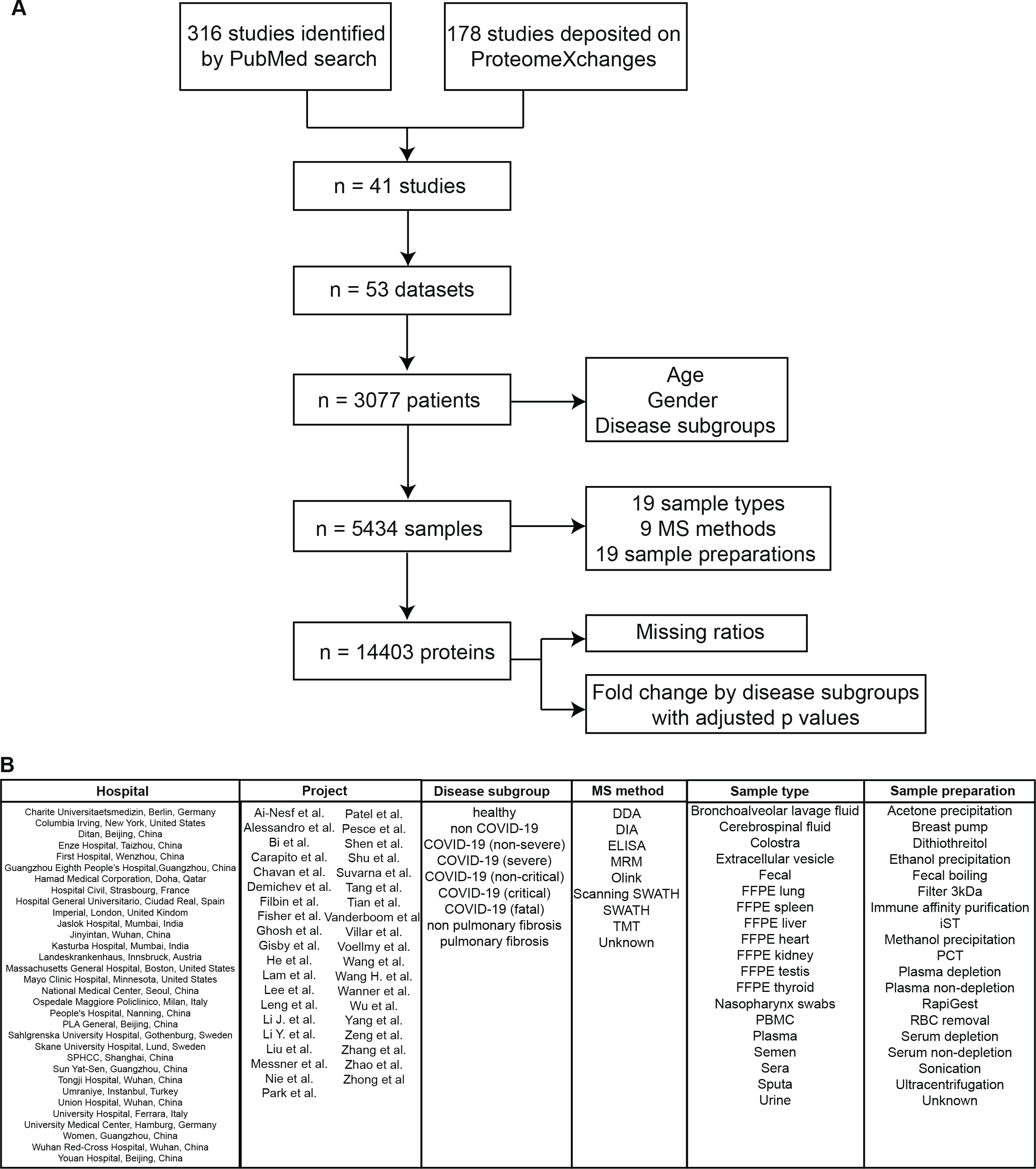
Study design. (A) Selection of a set of COVID-19 proteomics datasets. (B) List of the COVID-19 studies selected for our database.

Of the 1,794 patients with gender and age information, 60.4% were male and the median age was 49.1 years with a standard deviation of 17.12. Patients were categorized into seven disease subgroups. A total of 1083 COVID-19 patients had non-severe disease, while 629 had severe symptoms. Control cases, including healthy or non-COVID-19 patients, accounted for 19% and 12.5% of all cases, respectively. More than 80% of the samples were derived from blood: 50.4% from plasma and 30.3% from sera. Besides blood samples, urine samples constituted 9.6% and FFPE tissue 4.9% of all samples. As blood samples contain many high-abundance proteins that may interfere with the identification of low abundant ones, some studies performed additional depletion procedures on plasma or sera samples^56^. Specifically, 4.2% of the plasma and 28.9% of the serum samples were depleted of the highly abundant proteins. Regarding the mass spectrometry acquisition strategies used by the various studies, label-free quantification methods, including DDA (13.5%), DIA (7%), MRM (0.3%), scanning SWATH (2.5%), and SWATH (27.3%), were used for more than half of the samples. Otherwise, the Olink kit (27.3%) or TMT multiplexing methods (19.2%) were used (Table 1).

**Table 1.** Baseline characteristics of the patient included.

|  |  |
| --- | --- |
| **Gender** |  |
| Female | 745/1847(40.3%) |
| Male | 1102/1847(59.7%) |
| **Age** | 49.1(17.12)(n=1794) |
| **Disease subgroup** |  |
| Healthy | 586/3077(19%) |
| Non COVID-19 | 385/3077(12.5%) |
| COVID-19 (non-severe) | 1083/3077(35.2%) |
| COVID-19 (severe) | 629/3077(20.4%) |
| COVID-19 (fatal) | 85/3077(2.8%) |
| COVID-19 (critical) | 212/3077(6.9%) |
| Non-pulmonary fibrosis | 6/3077(0.2%) |
| Pulmonary fibrosis | 21/3077(0.7%) |
| COVID-19 (non-critical) | 21/3077(0.7%) |
| Control IL-6 | 16/3077(0.5%) |
| Low IL-6 | 18/3077(0.6%) |
| Medium IL-6 | 5/3077(0.2%) |
| High IL-6 | 10/3077(0.3%) |
| **Clinical sample type** |  |
| Bronchoalveolar lavage fluid | 9/5434(0.2%) |
| Cerebrospinal fluid | 8/5434(0.1%) |
| Colostrum | 6/5434(0.1%) |
| Extracellular vesicle | 23/5434(0.4%) |
| Fecal | 72/5434(1.3%) |
| Heart FFPE | 38/5434(0.7%) |
| Kidney FFPE | 61/5434(1.1%) |
| Liver FFPE | 52/5434(1%) |
| Lung FFPE | 37/5434(0.7%) |
| Nasopharynx swabs | 16/5434(0.3%) |
| PBMC | 103/5434(1.9%) |
| Plasma | 2737/5434(50.4%) |
| Semen | 27/5434(0.5%) |
| Sera | 1645/5434(30.3%) |
| Spleen FFPE | 32/5434(0.6%) |
| Sputa | 13/5434(0.2%) |
| Testis FFPE | 15/5434(0.3%) |
| Thyroid FFPE | 29/5434(0.5%) |
| Urine | 511/5434(9.4%) |
| **Sample preparation method** |  |
| Acetone precipitation | 149/5434(2.7%) |
| Breast pump | 6/5434(0.1%) |
| DTT | 44/5434(0.8%) |
| Ethanol precipitation | 27/5434(0.5%) |
| Fecal boiling | 72/5434(1.3%) |
| filter 3kDa | 10/5434(0.2%) |
| Immune affinity purification | 11/5434(0.2%) |
| iST kit | 77/5434(1.4%) |
| Methanol precipitation | 16/5434(0.3%) |
| PCT | 248/5434(4.6%) |
| Plasma depletion | 115/5434(2.1%) |
| Plasma non-depletion | 2535/5434(46.7%) |
| RapiGest | 13/5434(0.2%) |
| RBC removal | 8/5434(0.1%) |
| Serum depletion | 476/5434(8.8%) |
| Serum non-depletion | 1169/5434(21.5%) |
| Sonication | 16/5434(0.3%) |
| Ultracentrifugation | 329/5434(6.1%) |
| Unknown | 113/5434(2.1%) |
| **MS methods** |  |
| DDA | 732/5434(13.5%) |
| DIA | 381/5434(7%) |
| ELISA | 11/5434(0.2%) |
| MRM | 19/5434(0.3%) |
| Olink | 1491/5434(27.4%) |
| Scanning SWATH | 134/5434(2.5%) |
| SWATH | 1485/5434(27.3%) |
| TMT | 1044/5434(19.2%) |
| Unknown | 137/5434(2.5%) |

We next describe the analyses performed on the collected data and the results found. Specifically, after general data evaluations, we performed protein, pathway, and integrative analyses. We then demonstrate one of the possible use cases for the COVIDpro database.

To gain an overview of the various datasets, we first report the number of proteins identified by each study with the mass spectrometry’s missed detecting ratios. Although more than 14,403 proteins were measure by mass spectrometry across all projects, the largest proportion of proteins were identified in non-sera and non-plasma datasets. More than ten thousand proteins were detected in FFPE samples, while nasopharynx swabs accounted for over six thousand proteins, urine for about three thousand proteins, and several hundreds of proteins were measured in most sera and plasma samples. The number of proteins identified varies between different clinical specimens (Figure 2A). The number of patients and their disease subgroups are shown in Figure 2B. Most individual datasets described a few dozens to over a hundred patients; the only exception was a study that profiled 384 patients, which had high missing value rates.

**Figure 2.**
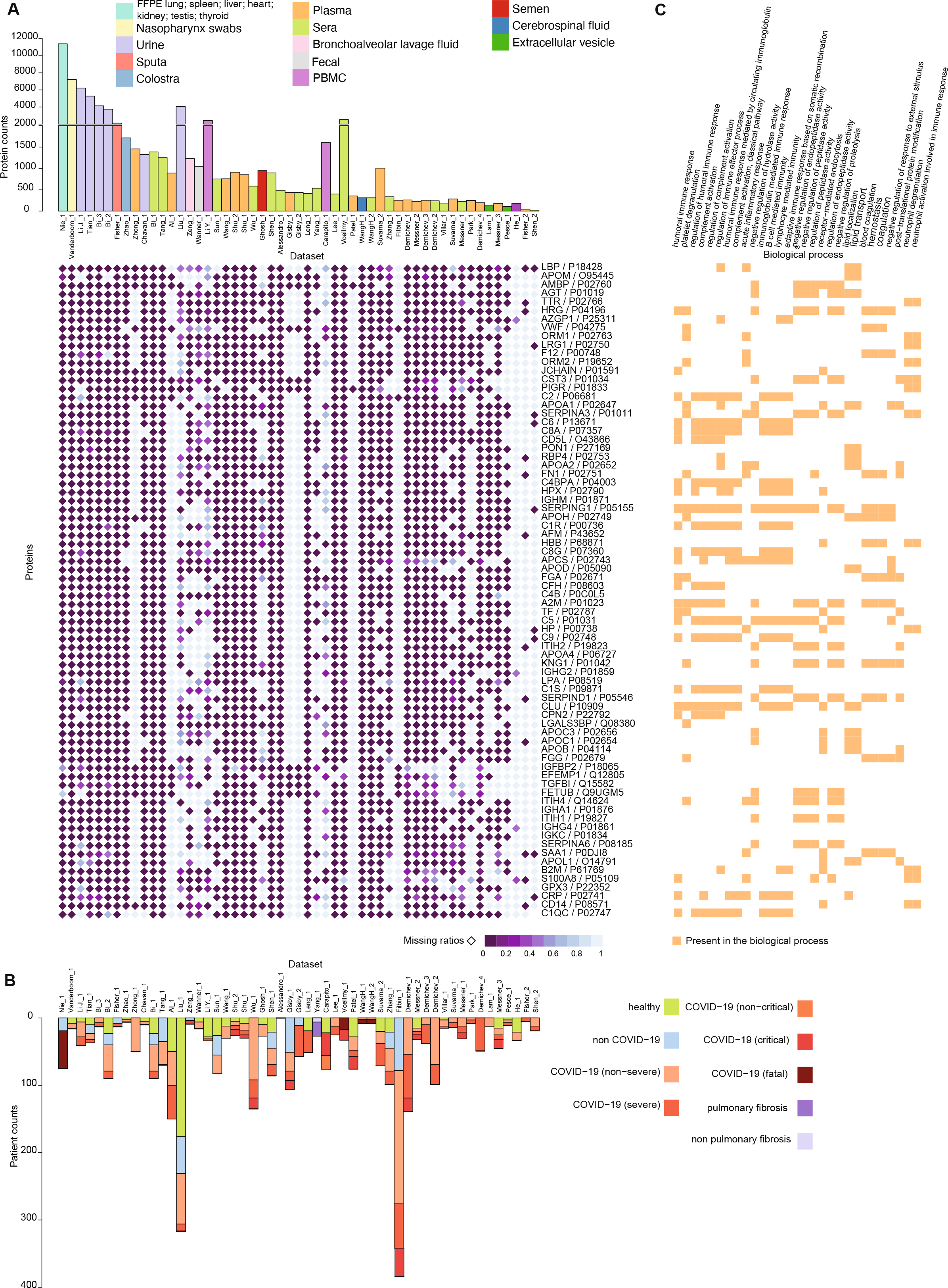
Protein expression in the COVID-19 datasets selected for COVIDpro. (A) Upper panel: the number of proteins in each dataset; lower panel: the 76 most frequently characterized proteins in the database. (B) Number of patients involved in each study. (C) GO enrichment of the biological processes involving the 76 most frequently identified proteins.

Next, to perform functional analyses, we focused on 66 proteins identified in at least 70% of the studies. Using a Gene Ontology (GO) analysis, we found that most of the identified proteins were involved in the immune response and the activation of the complement system (Figure 2C), which is consistent with previous findings that these proteins were more involved in the regulation of COVID-19 severity^9^. We then further evaluated the most frequently differentially expressed proteins. The most frequently appearing protein is the lipopolysaccharide binding protein (LBP) which binds to lipopolysaccharide (LPS). The latter has been reported to bind to SARS-CoV-2 S protein^57^. LBP is known to increase in the presence of bacterial infections and is a marker of sepsis^58–60^. It has been suggested that in COVID-19 patients this LBP increase is caused by dysfunction of the gut-blood barrier that leads to increased microbial translocation^61,62^

We then compared the expression of LBP across all the studies where it was detected. The level of LBP increased significantly with the severity of disease, from healthy to non-severe and severe groups, when sera and plasma samples were analyzed. By contrast, we observed different LBP dynamics in urine, EV, colostrum, and cerebrospinal fluid samples (Figure 3A). In COVID-19 autopsies, LBP was seen to have significantly decreased in the kidneys and lungs (Figure 3B). Also, LBP showed slightly different expression dynamics in COVID-19 patients with prolonged RNA shedding (Figure 3C). Furthermore, the level of IL-6 also positively correlated with the level of LBP (Figure 3D); IL-6 is known to be involved in both fever and inflammation responses^63,64^. The expression of IL-6 decreased in plasma during convalescence (Figure 3E). Finally, in extracellular vesicle samples, LBP increased and then decreased around the third week (Figure 3F). As elevated levels of LBP have also been observed in infections and inflammatory diseases, this protein could be an indicator for the severity progression of COVID-19.

**Figure 3.**
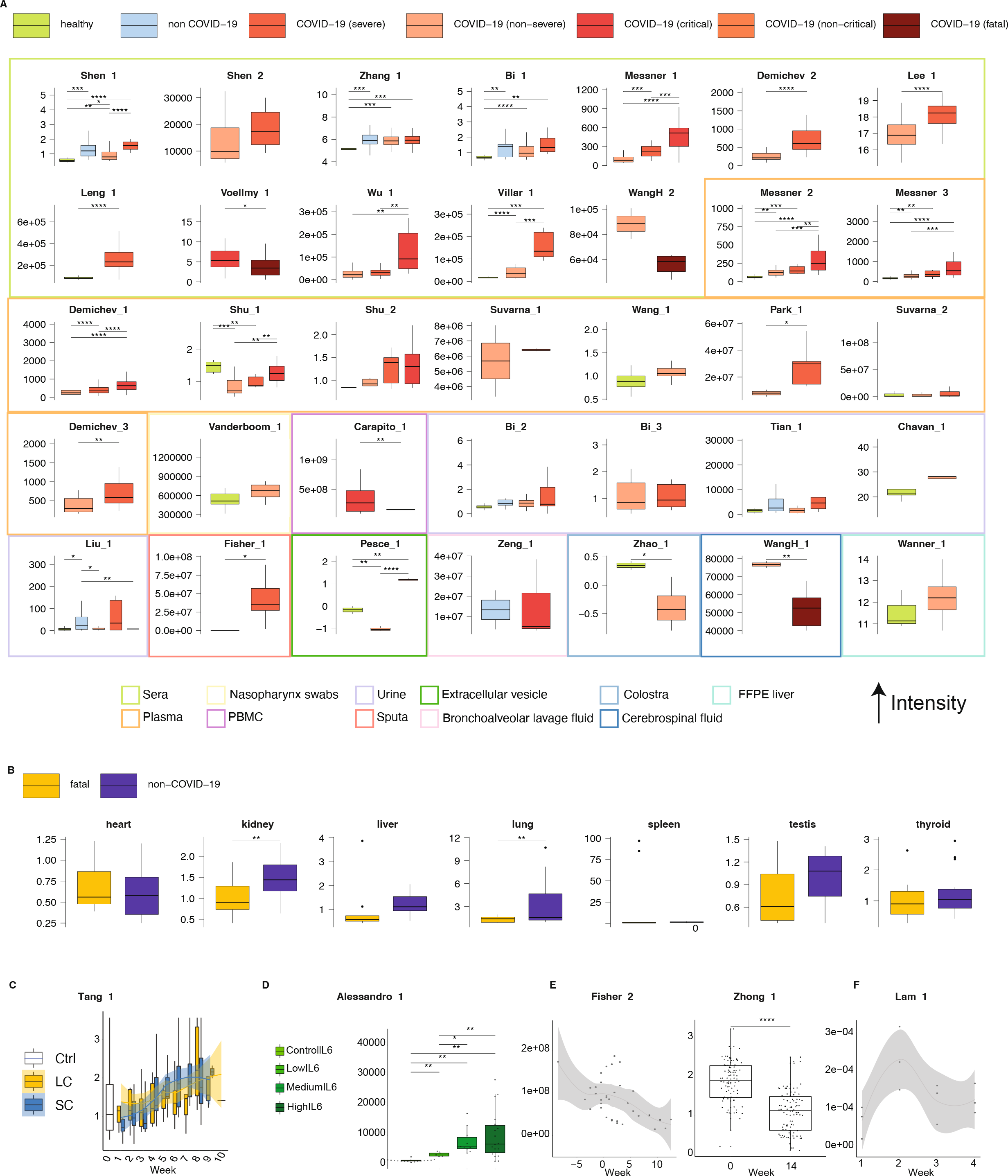
Meta-analysis of LBP expression. (A) Expression of LBP in non-longitudinal studies. (B) Expression of LBP in seven FFPE tissues. (C) Expression of LBP in a study including cases with either a long (LC) or a short course (SC) of the disease. (D) Expression of LBP in a study where samples were grouped according to IL-6 expression. (E) Expression of LBP in two longitudinal plasma studies. (F) Expression of LBP in longitudinal EV samples.

Next, we analyzed and compared the urine proteomes of non-severe and healthy patients from the Bi_2 dataset using the Student's *t*-test. We found that 59 and 839 proteins were up- and down-regulated, respectively, in severe patients (Figure 4A). Using GO (Figure 4B) and KEGG enrichment analyses (Figure 4C), we discovered that a large proportion of the up-regulated proteins were involved in the central carbon cycles, while the down-regulated ones were associated with binding and adhesion proteins. Empowered by cyjshiny^55^ package and KEGG pathway interactions taken from Pathway Commons version 12^65^, which contains 79 common metabolic pathways, we identified 74 pathways containing these dysregulated proteins. Six pathways having higher number of differentially changed proteins are shown in Figure 4D. The down-regulated proteins were illustrated in the metabolism of the amino acids while most up-regulated proteins were involved in the TCA cycles (Figure 4D). Compared with the non-severe patients, many proteins involved in glycolysis and glucogenesis were down-regulated in the urine samples of the severe cases; however, only a few proteins involved in glycolysis and glucogenesis were dysregulated in the blood samples (Figure 4E).

**Figure 4.**
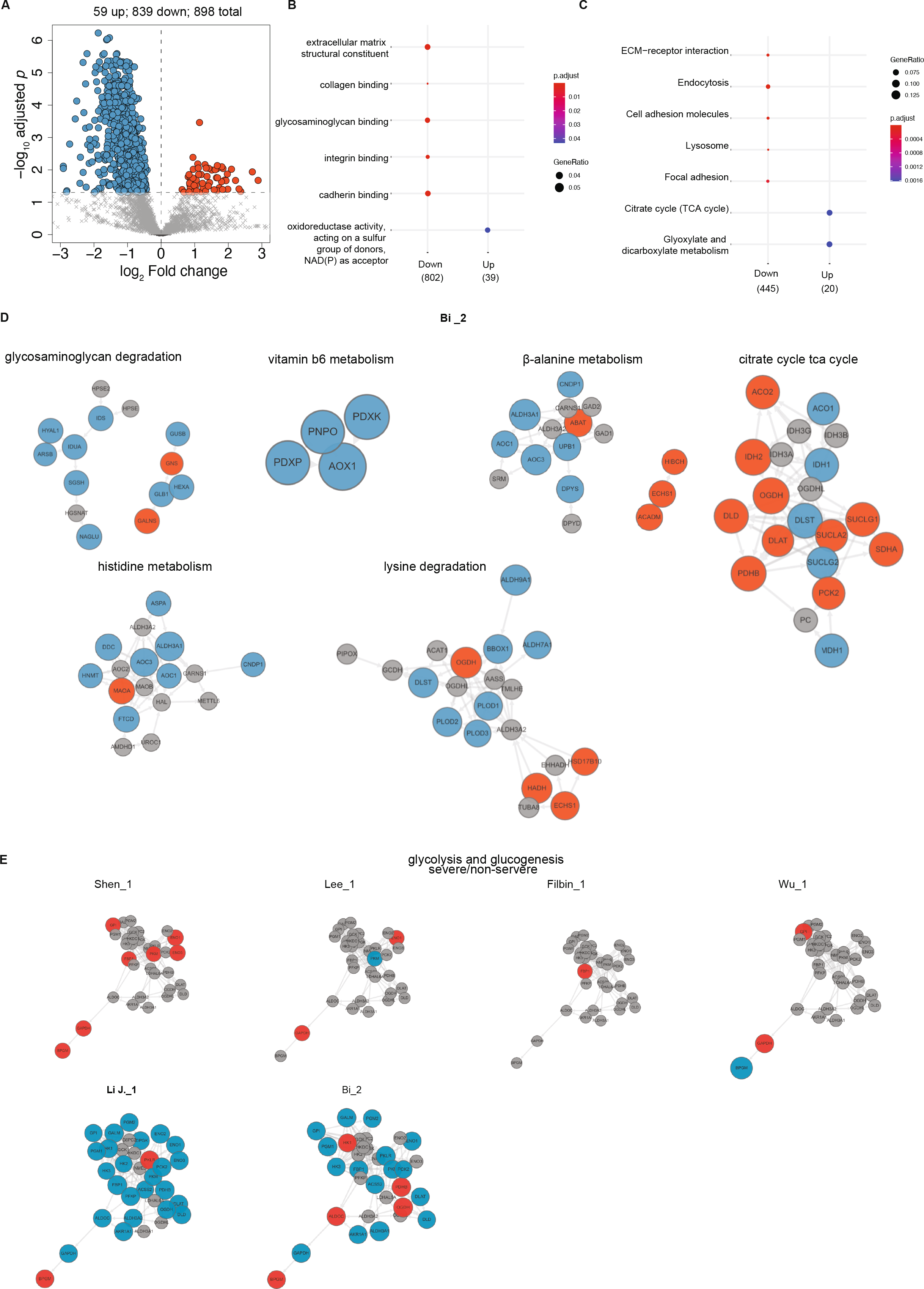
Pathway analysis of the differentially expressed proteins of COVID-19 patients. (A) Volcano plots for differentially expressed proteins (DEPs) in non-severe vs. healthy cases from the Bi_2 dataset. (B) GO enrichment results of the up- and down-regulated DEPs. (C) KEGG enrichment results of the up- and down-regulated DEPs. (D) Selected pathways from the Bi_2 dataset involving dysregulated proteins. Red (up) and blue (down) indicate the direction of the regulation. (E) Differentially changed proteins in glycolysis and glucogenesis for severe and non-severe cases.

COVIDpro’s datasets can be used to explore and validate diagnostic biomarkers of COVID-19. As a proof of principle, we generated a machine learning model for classifying COVID-19 severity (severe vs. non-severe) based on specific proteins. First, using the tools provided in our server, we identified the differentially expressed proteins between severe and non-severe patients across all the studies that included these two categories (Figure 5A). We thus focused on 51 differentially expressed proteins that appeared in at least five studies. Next, we built a preliminary random forest model to classify COVID-19 severity using the Shen_1 dataset as the training set (Figure 5B). The top nine proteins were selected: SAA1, SERPINA1, angiotensinogen (AGT), C9, LRG1, HABP2, SERPINA3, HRG, and HP (Figure 5C). These nine proteins were used to build a random forest-based classifier, which correctly classified all COVID-19 cases from the Shen_1 dataset. Our classifier was then further validated using five independent datasets, achieving a mean area under the curve (AUC) of 0.87 and a mean accuracy (ACC) of 0.79 (Figure 5D). Many of the selected proteins, including the acute phase proteins SAA1 as well as the complement activation protein C9, have been associated with severe patients^66^. In addition, the serine protease inhibitor SERPINA1/3 has been reported to inhibit the viral spike protein TRMPRSS2^67^. AGT was also a selected model feature. The enzymatic product of AGT is the precursor of angiotensin II, which is the substrate of the host protein angiotensin-converting enzyme (ACE) homolog-2 (ACE2). As a consequence, severe diseased COVID-19 patients have exhibited an elevated expression of AGT^68^. Of the genes we identified as part of our model, HABP2 has been studied less thoroughly in connection with COVID-19, and our work suggests it warrants further study. HABP2 plays a role in blood coagulation and may be involved in the abnormalities of coagulation seen in COVID-19 patients^69,70^.

**Figure 5.**
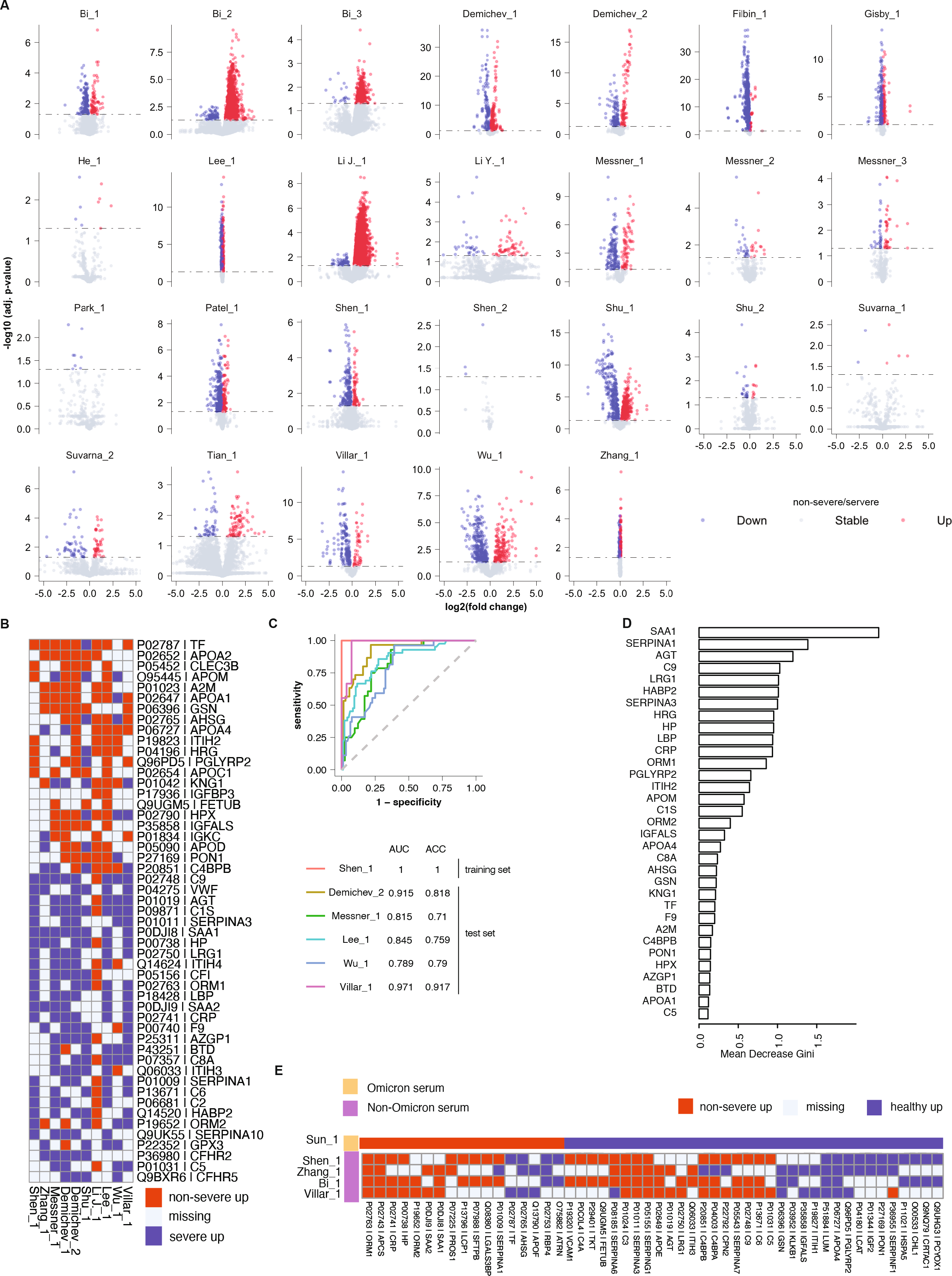
Differentially expressed proteins between severe and non-severe patients and machine learning modeling. (A) Volcano plots of the differentially expressed proteins in non-severe vs. severe patients across all the datasets containing these patient groups. (B) The 51 dysregulated proteins that appeared in at least five projects. (C) The nine features with the highest mean decrease Gini from the random forest model. (D) Performance of random forest classifier for training and independent validation cohorts, including Receiver operation curves (ROCs), Area under curves (AUCs), and accuracies (ACCs).

## Discussion

In this study, we generated a large public database, COVIDpro, including the most relevant published proteomics datasets of COVID-19 patients. We also showed the results of a set of analyses we performed using the toolkits available in COVIDpro. The COVIDpro database covered the published proteomics data of COVID-19 patients till May 2022, containing 3077 patient cases, 5434 samples from 19 of sample types, and 14,403 proteins profiled. This data resource allows performing meta-analyses of protein regulations across multiple clinical specimens of COVID-19 patients from eleven nations. For each protein from the 14,403 proteins included in COVIDpro, we developed a user-friendly interface for browsing its expression and pathway involvement across multiple datasets. This resource could be used to support biomarker and therapeutic discoveries for COVID-19. As a showcase, we used COVIDpro to identify biomarkers of COVID-19 severity and perform *in silico* validation experiments. To the best of our knowledge, this is the most comprehensive COVID-19 protein expression repository.

We included a module to search for the latest relevant literature and another to append new datasets to this database resource, allowing its timely update. The sever will be maintained every quarter in the coming few years. The collected proteomic datasets were downloadable as readable text tables or in an R object RDS format, allowing other researchers to re-analyze the data for new discoveries. For example, identifications of clusters of proteins that go beyond the severity of the disease; dysregulated proteins in different ages, genders and geographical locations; specific patterns in the immune response for vaccine development.

Our data-driven study was different from the hypothesis-driven research, where more combinations of results could be shown depending the questions to address using our online database application. Here we only show one typical result and its interpretation due to the space limitation. In addition, our study is phenomenological by nature for the observance of the measured data, molecular functional validation cannot be surrogated to confirm the dysregulated proteins as therapeutic targets or potential biomarkers for diagnostic prediction.

Constructed with an R shiny framework, the COVIDpro analysis pipeline works as a cross-platform browser application that does not require any software installation. The R shiny framework integrates well with JavaScript and Cascading Style Sheets (CSS), allowing customized analysis modules to be generated. Our easy-to-access application allows users to explore COVID-19 proteomics datasets and validate their hypotheses. This COVID-19pro database may be a useful resource for nagivating dysregulated proteins in various clinical specimens from patients with COVID-19.

